# Rapid learning of the 5-choice serial reaction time task in an automated rodent training system

**DOI:** 10.1101/2020.02.16.951491

**Authors:** Eszter Birtalan, Anita Bánhidi, Joshua I. Sanders, Diána Balázsfi, Balázs Hangya

## Abstract

Experiments aiming to understand sensory-motor systems, cognition and behavior often require animals trained to perform complex tasks. Traditional training protocols require lab personnel to move the animals between home cages and training chambers, to start and end training sessions, and in some cases, to hand-control each training trial. Human labor not only limits the amount of training per day, but also introduces several sources of variability and may increase animal stress. Here we present an automated training system for the 5-choice serial reaction time task (5CSRTT), a classic rodent task often used to test sensory detection, sustained attention and impulsivity. We found that fully automated training without human intervention greatly increased the speed and efficiency of learning, and decreased stress as measured by corticosterone levels. Introducing training breaks did not cancel these beneficial effects of automated training, and mice readily generalized across training systems when transferred from automated to manual protocols. Additionally, we validated our automated training system with mice implanted with wireless optogenetic stimulators, expanding the breadth of experimental needs our system may fulfill. Our automated 5CSRTT system can serve as a prototype for fully automated behavioral training, with methods and principles transferrable to a range of rodent tasks.

## Introduction

In behavioral neuroscience, animal training requires a costly investment of work hours and resources. It is a major undertaking requiring human accuracy and persistence, constraining efforts to standardize and scale up behavioral experiments. There is an increasing need for high-throughput behavioral assays as systems neuroscience moves towards increasingly more complex behaviors, optogenetic manipulations and recording neural activity via electrophysiology or imaging in behaving animals^1^.

Systematic studies found that uncontrolled factors may have profound impact on the experimental results^2–4^. Moreover, potential subconscious biases of the experimenters may pose even larger problems than serendipitous differences. This is especially important in pharmacology and optogenetic experiments, where different handling of the treated and control groups, even in subtle ways, may introduce false positive results. Blinding the experimenter to the group identities averages such differences out as a consequence of the strong law of large numbers^5,6^; however, blinding is often not possible due to overt differences between experimental groups and such convergence of the mean to the expected value may take prohibitively large samples^7^.

A few automated training systems have been developed for rodent behavioral tasks^8–15^, including 5-choice serial reaction time task (5CSRTT)^16,17^, in order to standardize the training and reduce the effects of human factors and other random variables. While these systems provide means for large capacity automated training of rodents, most of them are customized to train a specific task variant, and/or contain expensive, proprietary components. For these reasons, automated behavioral training of the 5CSRTT task has not yet become widespread. Here we developed an affordable, open source, high-throughput automated training system for mice and demonstrate its use on an automated protocol of the widely used 5CSRTT assay^18–21^. We show that use of this Automated Training System (ATS) allows faster training of mice, and that improved training time results from the higher number of trials performed daily. To improve upon existing systems described in literature, we (i) provide an inexpensive, modular, open source training setup, (ii) fully eliminate human interaction with the animals during training, (iii) evaluate the effects of training breaks and transfer from automated to manual training setups, (iv) demonstrate that automated training reduces stress compared to traditional training and (v) validate use of our training setup with wireless optogenetics to increase the range of possible experiments the assay is capable of.

## Results

### Stable performance despite decreased activity in the afternoon (middle of the light phase)

We developed a fully automated, open source, modular training system, in which a training chamber was connected to two separate home cages, each housing a single mouse. Access to the training chamber was controlled by motorized gates, and mice were allowed to enter the training chamber based on a fixed, regular schedule of 15 minutes training every two hours (Fig. 1-2; Methods).

**Figure 1.**
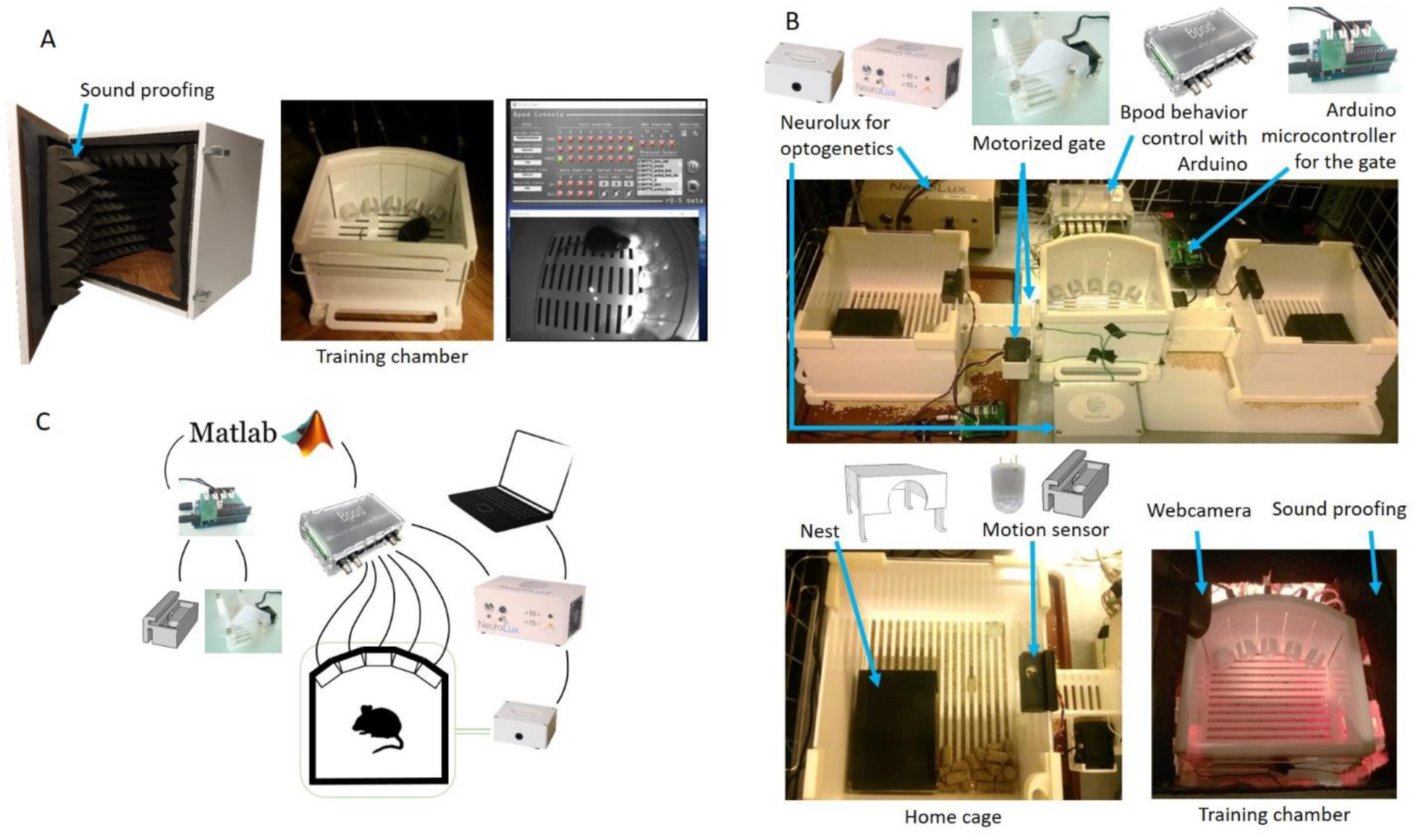
Behavioral setup. (A) Manual training setup. Left, the training chamber was placed in a sound attenuated wooden box (60×60×60 cm). Middle, the training chamber housed five water ports (Sanworks) with infrared sensors and LEDs. Right, the water ports were controlled by the Bpod behavior control unit (Sanworks) during training (top), while the animal was monitored via a high definition camera (FlyCapture; bottom). (B) Automated training setup. The ATS (top) consisted of a training chamber (bottom right) identical to that of the manually trained animals except for the side openings, through which it was connected to home cages (bottom left) on both sides. The home cages were equipped with a nest for the animals and a motion sensor (Panasonic) attached to the roof. The home cages were connected to the training chamber via tunnels blocked by motorized gates controlled by an Arduino. The equipment for wireless optogenetics (Neurolux) and the Bpod behavior control unit were placed outside the ATS. (C) Schematic of the hardware-software connections of the ATS and wireless optogenetics. The Neurolux control unit and the water ports were connected to Bpod, whereas the motorized gate and motion sensors were connected to their corresponding Arduinos. The Bpod and Arduinos were connected to the computer and controlled by the same Matlab code (available at https://github.com/sanworks/Pipeline_Gate and https://github.com/hangyabalazs/ATS).

**Figure 2.**
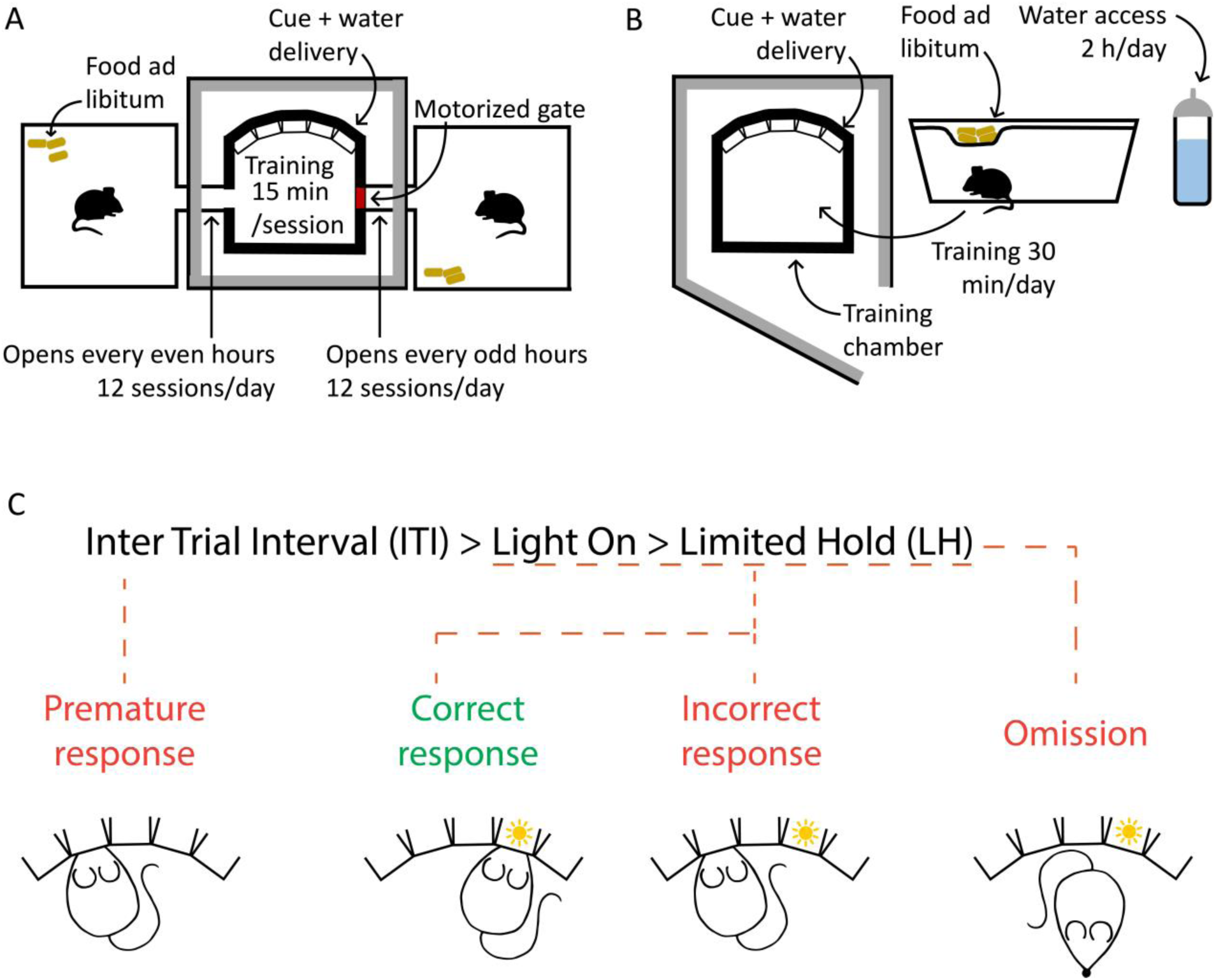
Training protocol. (A) Schematics of ATS training. All animals had access to food *ad libitum* in their home cages, whereas they received water in the training chamber, accessible for 15 minutes in every two hours (free access to water at the beginning of each session and water rewards during training). (B) Schematics of manual training. Animals were kept in standard mouse cages with access to food *ad libitum.* Water was freely available for two hours/day. Mice were moved to the training chamber for 30 minutes training sessions daily, where they received additional water as reward, then moved back to their home cages. (C) Trial phases and possible outcomes of the 5-choice serial reaction time task (see details in ref.^20^).

A group of 12 mice were trained on a 5CSRTT in the ATS (see Methods). Every two hours, an open gate gave mice the option to enter the training chamber or skip a session. This allowed us to test whether mice show a natural preference towards particular times of the day for training and whether accuracy in the 5CSRTT depended on what time the session was performed. The mice were kept on 12-hour light/dark cycle, with light phase starting at 7 am. We found that mice were least active between 3 and 4 pm, showing significantly lower probability of entering the training chamber (entry probability 3-4 pm, mean ± SEM, 0.45 ± 0.08; p < 0.05 compared to 1-10 am and 5-12 pm, Fig. 3) and more omissions during training (mean ± SEM, 20.93 ± 4.3%, p < 0.05 compared to 23 pm-4 am and 11-14 am). Entry probability gradually declined from 9 am to 4 pm, then steeply increased to reach a maximum of 0.92 ± 0.03 (mean ± SEM) in the last hour of the day. While entry probability varied with circadian time, accuracy did not show significant fluctuations throughout the day (Fig. 3).

**Figure 3.**
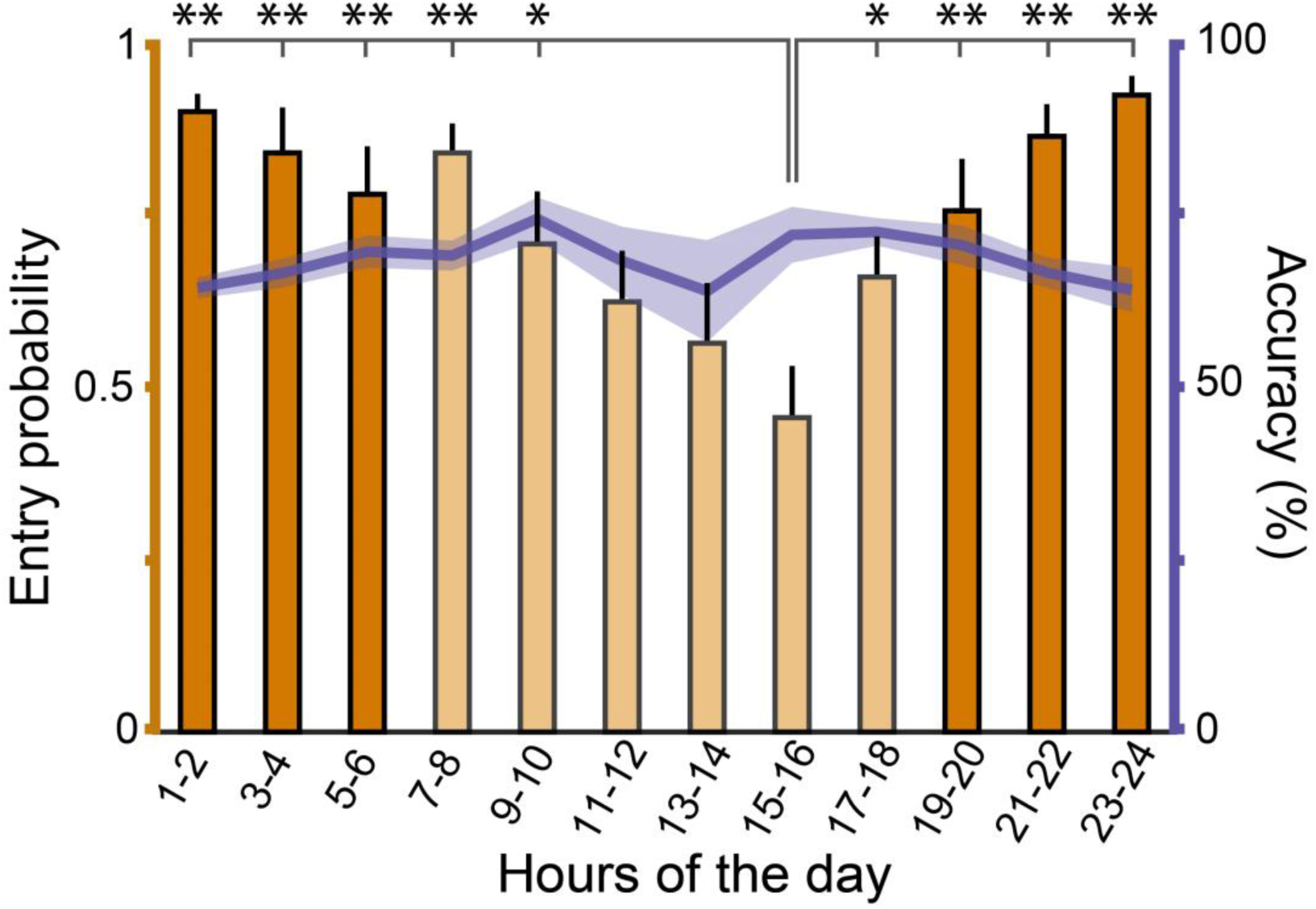
Dependence of activity and performance of ATS-trained mice on the time of day. Activity (bar graphs, y axis is on the left) was defined as the probability of mice engaging in a training session (sessions performed / number of available sessions). Light phase (indicated by lighter colors) started at 7 am. The animals’ accuracy (line plot, y axis is on the right) was stable during the day. Bars and line plot show mean ± SEM. *, p < 0.05, ** p < 0.01, t-test; N = 12.

### Mice learn faster in the ATS compared to traditional manual training

To evaluate performance of mice in the custom-developed ATS, ATS-trained mice were compared to a cohort of mice (N = 14) trained manually by expert personnel. Manual training was carried out according to Bari et al.^20,22^ in single daily sessions between 9 am and 12 pm and lasted approximately 30 minutes (see Methods). Additionally, to test if stereotaxic surgery and implantation had any effect on the performance of the animals, a third group of mice (N = 7), implanted with head-mounted LEDs for wireless optogenetics was trained in the ATS. These mice had been injected with control virus and were photostimulated during the inter-trial interval in 50% of the sessions (see Methods).

Learning performance was compared after one week of training (Fig. 4). Specifically, the average of a theoretical maximum of 12 sessions in the ATS on day 7 was compared to the single manual training session on the corresponding day in the traditional setup. Mice advanced through the twelve classical training stages of 5CSRTT defined by Bari et al.^20^ automatically based on their performance; therefore, it was possible to compare the training stages they reached by the end of one week. Half of ATS-trained animals reached the highest, twelfth stage, and all of them advanced beyond stage 5. In contrast, manually trained animals did not pass the third stage by the end of the week, achieved by 71% of the animals. Thus, we found that mice learned significantly faster in the ATS (Fig. 4A, F_2,30_ = 73.29, p < 0.0001; one-way ANOVA). Implanted mice reached slightly but significantly lower levels than non-implanted mice in the ATS (p < 0.05), while they were substantially more advanced than manually trained mice (p < 0.001, Newman-Keuls post-hoc test).

**Figure 4.**
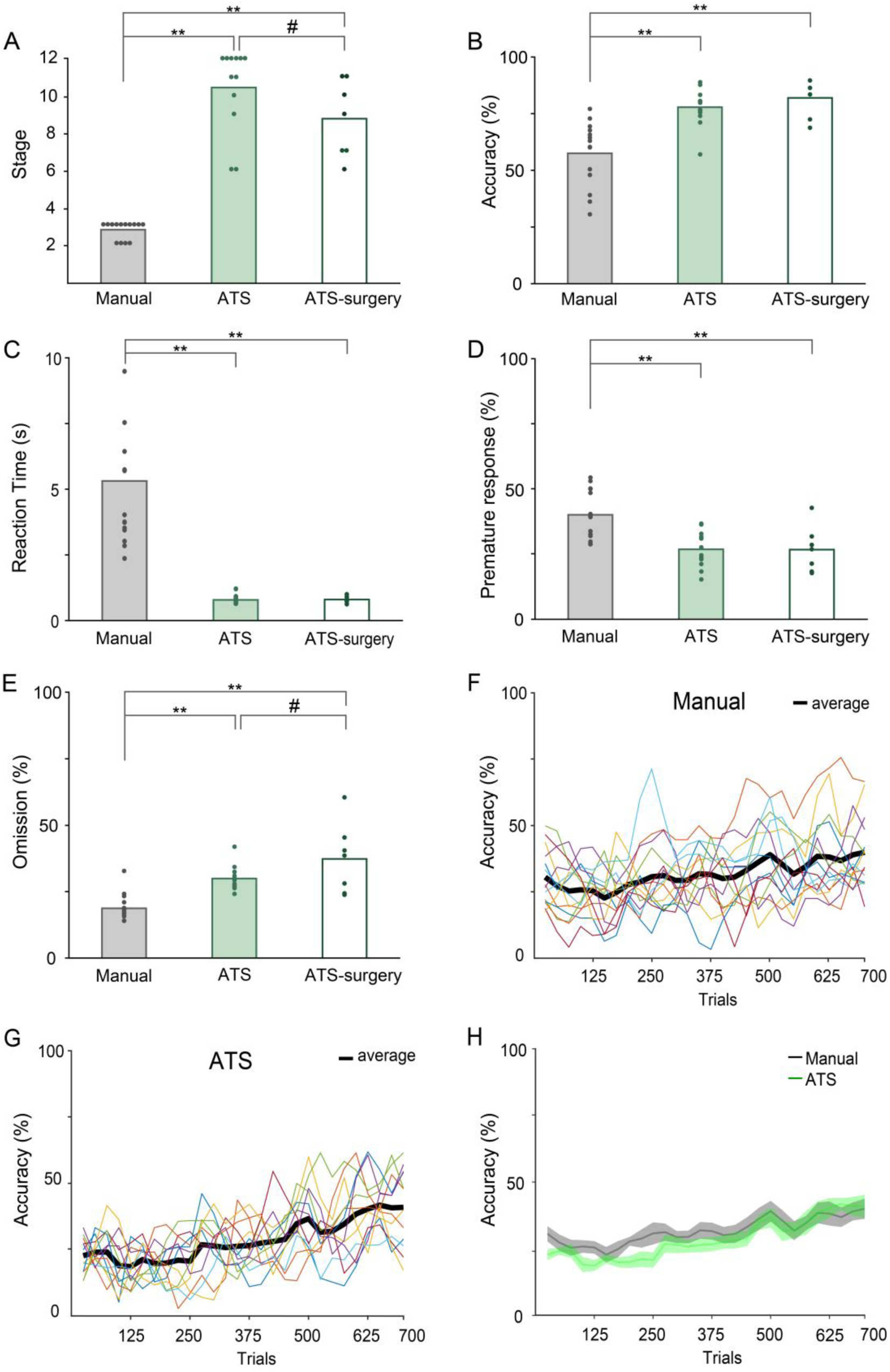
Comparison of one week of manual and ATS training, with and without surgery. (A-E) Performance during the 7th day of training compared between groups. Bar, mean; dots, individual mice. Mice trained in the ATS reached higher stages (A), performed with higher accuracy (B) and shorter reaction times (C). They performed fewer premature responses (D) but omitted more trials (E). (F-G) Accuracy calculated for the first 700 trials of training (in 50 trial-windows with 50% overlap) for manually (F) and ATS-trained (G) mice. Colored lines, individual mice; black line, average. (H) Average accuracy in the first 700 trials in the manual (grey) and ATS group (green); lines and error shades represent mean ± SEM. *p < 0.5, ** p < 0.01; one-way ANOVA (A-E) and repeated-measures ANOVA (H); manual, N = 14; ATS, N = 12; ATS-surgery, N = 7.

Beyond reaching higher stages in the ATS, we found significant main effects between the three groups in all performance measures tested (accuracy, F_2,30_ = 15.34, p < 0.0001; reaction time, F_2,30_ = 21.88, p < 0.0001; premature responses, F_2,30_ = 10.26, p < 0.001; omissions, F_2,30_ = 16.34, p < 0.0001; one-way ANOVA, Fig. 4B-E). Post-hoc tests revealed that ATS-trained mice were significantly more accurate than manually trained animals, regardless whether implantation surgery was performed before the ATS training (intact ATS, p < 0.001; implanted ATS, p < 0.001; Fig. 4B). No significant difference in accuracy between the implanted and intact mice trained in ATS was found (p = 0.428).

While the time windows in which mouse responses to cue stimuli were accepted varied across training stages, all mice had at least 5 seconds to perform a correct response. Mice trained in the ATS typically performed fast responses (mean ± SEM, 0.79 ± 0.05) with significantly shorter reaction time than manually trained animals (mean ± SEM, 5.31 ± 0.79; p < 0.001, Fig. 4C). Implantation surgery did not lead to a difference in reaction times (p = 0.999). We also found that ATS-trained mice performed less premature responses (p < 0.01) but omitted more trials (p < 0.01) than the manually trained animals (Fig. 4D-E). Implanted mice omitted more trials than intact animals in the ATS (p < 0.05, Fig.4E).

To dissociate whether better performance of ATS-trained animals was due to a steeper learning curve, higher number of trials performed (ATS, mean ± SEM, 741 ± 23 trials/day; manual, mean ± SEM, 143 ± 10 trials/day) or a combination of both, we compared performance improvement in the two training groups for the first 700 trials completed, calculated in 50-trial sliding windows (50% overlap; Fig. 4F-H). We found similar learning curves (group, F_1,24_ = 1.75, p = 0.20; time, F_28,672_ = 11.95, p < 0.0001; time x group, F_28,672_ = 1.23, p = 0.19) in the two groups when plotted as a function of completed trials, suggesting that the ATS-trained animals showed an increased performance compared to traditional manual training due to the large number of trials mice completed during the 12 possible daily sessions ().

### The benefits of the ATS are not cancelled by training breaks

Optimal design of electrophysiology or optogenetics experiments often requires a training period, followed by surgery and recovery, after which training is resumed, combined with recording or manipulating a selected set of neurons. Typically, this leads to a transient drop in performance – so we sought to determine whether such a protocol would cancel some of the benefits of the ATS.

Therefore, we measured the efficiency of both manual and ATS training interrupted by pauses (Fig. 5A). First, a one-week training period was performed as shown previously (Fig. 4), then a 17-days pause was introduced to model training breaks introduced by surgery and recovery (manual, N = 8; ATS, N = 4 mice). After the pause, training was resumed from the stage mice had reached by the end of the first week of training period. Compared to day 7, ATS-trained mice showed a transient decrease in accuracy after the pause (Fig. 5C; p = 0.06, Wilcoxon signed rank test between accuracy at day 7 and 25 in the ATS; larger accuracy change after the pause for ATS vs. manual training, p < 0.05, Mann-Whitney U-test) that vanished after an additional week of training (day 31), reaching pre-pause levels. Note however, that manually trained animals only reached stage 2 on average by day 7, thus resumed training at an earlier training stage compared to ATS-trained mice, trained at stage 8 on average (Fig. 5B; p < 0.01, time × training group interaction, repeated-measures ANOVA).

**Figure 5.**
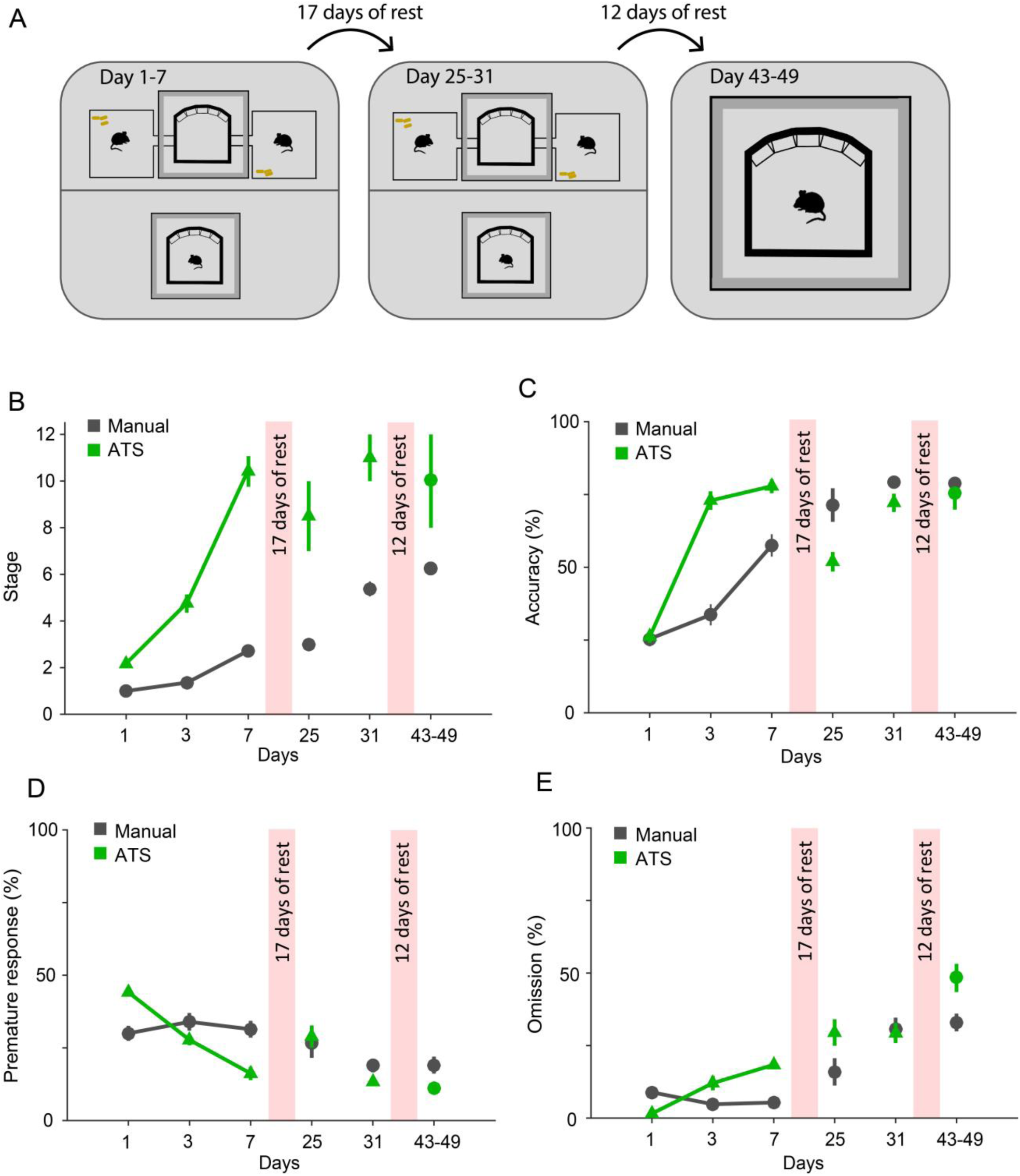
Effect of training breaks on performance. (A) Schematics of the experiment. One week of training was followed by a 17-days-long training break, after which mice resumed training from their previous stages in the same setup for another week. Following a second break of 12 days, all mice were transferred to the manual training setup. (B-E) Comparison of stage (B), accuracy (C), premature responses (D) and omissions (E) between the manual and ATS groups. After the first break, ATS-trained animals’ accuracy decreased compared to the manually trained group, which difference disappeared after one week of training. We did not find a significant difference in the studied parameters after the second break and transfer to manual training. All values represent mean ± SEM.

Proportion of omissions increased, while premature responses decreased throughout the training weeks for both ATS and manually trained mice (Fig. 5D-E; omission, F_4,32_ = 9.56, p < 0.0001; premature response, F_4,32_ = 5.98, p < 0.01; repeated-measures ANOVA). The ratio of omissions and premature responses was not significantly affected by the training break (p > 0.05, Wilcoxon signed rank test for within group and Mann-Whitney U-test for between group comparisons).

In practice, electrophysiology experiments may require large implants, head stages and tethering of mice to data acquisition equipment during the behavioral experiments, precluding use of ATS. Nevertheless, ATS may still speed up such experiments by allowing rapid pretraining of mice before implantation. However, in this case mice are switched from ATS to manual training, which may lead to significant drop in performance if mice fail to generalize over the training systems. To address this, we introduced a 12-day-long second break of training with the same mice (manual, N = 2; ATS, N = 2 mice), after which ATS-trained mice were transferred to the manual training setup (Fig. 5A-E). This second pause (from day 31 to day 43) and change of training protocols did not lead to performance drops in mice originally trained in the ATS, and demonstrated that a seamless transfer to manual training is possible while retaining the performance benefits of pretraining in ATS.

### Training in the ATS causes less stress for the animals

We hypothesized that ATS may cause less stress to mice, since they are not handled or in any other way disturbed by lab personnel, and are free to decide whether to engage in the training at every scheduled opportunity^23–25^. To test this, we collected blood samples and measured changes in the concentration of corticosterone, the main glucocorticoid hormone regulator of stress responses in rodents^26–29^. After the last behavioral session on the 7^th^ day of training between 9 am and 12 pm, mice were allowed (ATS-trained) or transferred (manually trained) to their home cages for 10 minutes, after which mice were transferred to a separate room for decapitation and blood sample collection (see Methods). Mice consumed comparable amounts of water in the ATS and manual setups before hormone testing. We found a significant main effect of corticosterone levels between groups (F_2,15_ = 22.81,p < 0.0001, Fig. 6A). Post hoc tests revealed that corticosterone concentration of the manually trained mice (N = 6) was significantly higher than that of the control (N = 6) and the ATS-trained groups (N = 6, p < 0.001 for both comparisons), while the ATS-trained group did not show a significant difference from the control group (p = 0.27, Fig. 6A). These results demonstrate that automated training causes less stress to mice compared to manual training and handling, despite the larger number of sessions, more completed trials and longer cumulative training time in the ATS.

**Figure 6.**
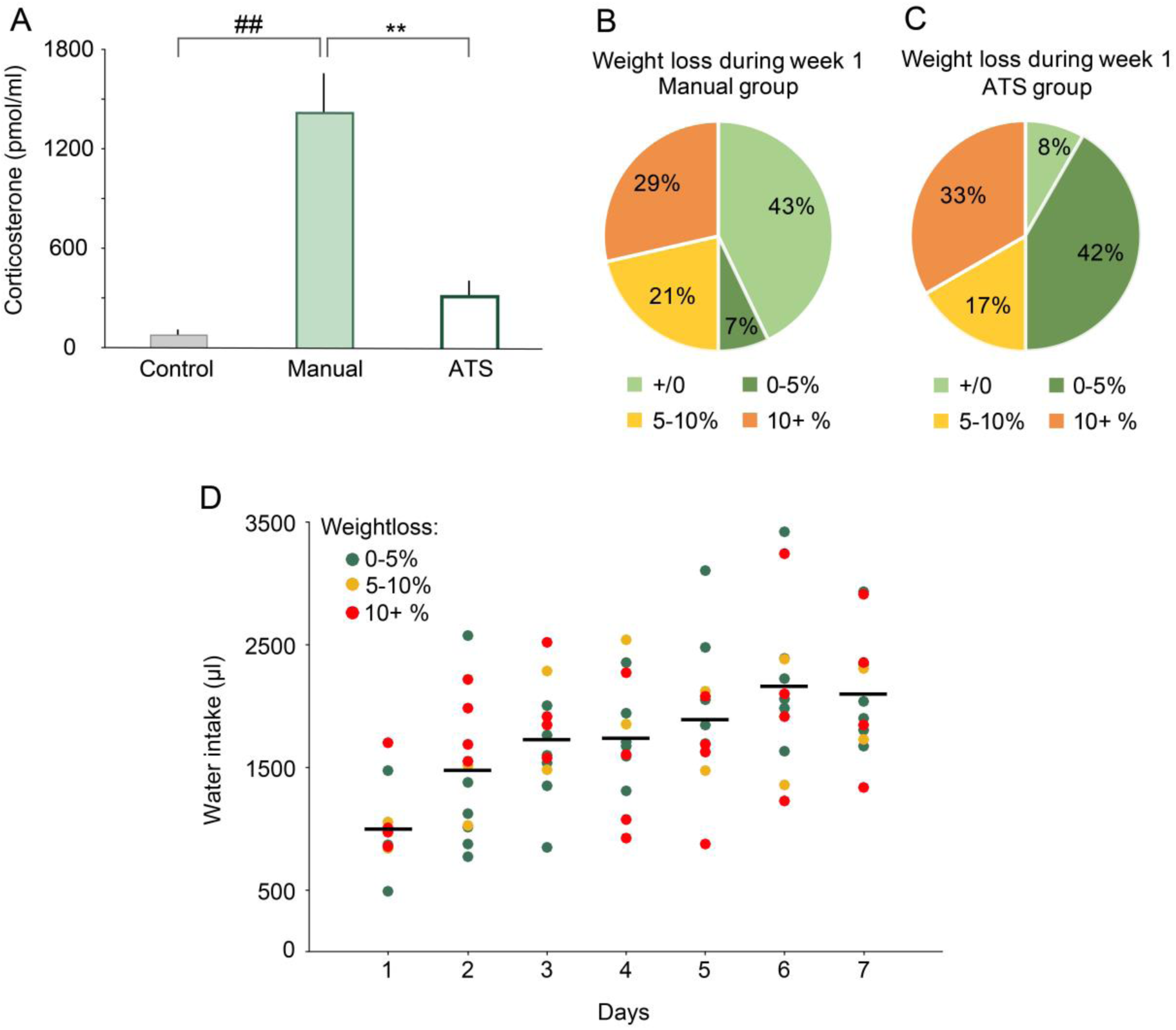
The effects of manual and ATS training protocols on stress hormone levels, bodyweight and water intake. (A) Blood corticosterone levels were higher in the manually trained mice when compared to the control or the ATS trained mice. There was no difference in blood corticosterone levels when comparing the control and the ATS trained mice. (B-C) Changes in body weight between training day 1 and 7 in the manually and ATS-trained mice. In both groups, 50% of the animals lost less than 5% of their bodyweight (this includes animals that gained weight). (D) There was no correlation between bodyweight change and water intake in the ATS-trained mice. Dots represent the water intake of individual mice color-coded according to their bodyweight-change after one week of training. Lines represent average water intake. ** p < 0.01, one-way ANOVA; control, N = 6; manual, N = 6; ATS, N = 6.

Finally, we monitored the weight of mice during training. Water restricted mice typically show a mild weight loss after the first week of training. We did not find a significant difference between weight changes in the ATS compared with manual training (F_1,10_ = 1.39, p = 0.27), although more animals tended to show mild weight loss in the ATS (Fig.6B-C). Surprisingly, weight changes did not show an obvious correlation with the cumulative water intake of the animals (p > 0.05, R = 0.117).

## Discussion

Rodents are capable of performing a large variety of cognitive tasks, which has rendered them a popular model for investigating how brain controls behavior. However, rodents have almost exclusively been trained manually by human trainers, which limits training efficiency and may introduce covert biases. Here we presented a fully automated training system (ATS) for 5-choice serial reaction time task, popular for investigating sensory detection, sustained attention and impulsivity^16,18,19,21,22^. Mice engaged in training voluntarily on a regular schedule without any human interference throughout the entire training period. We showed that training in the automated system was substantially faster and caused less stress to the animals. We equipped the training setup with wireless optogenetic stimulation. The ATS is modular, affordable, open source and can easily be adopted to a wide range of tasks.

Manual training on 5CSRTT may take 30-60 days or longer^30,31^. In contrast, we found that mice can be fully trained on 5CSRTT in the ATS in one weeks’ time. Half of the auto-trained animals reached the highest stage 12 according to Bari’s training protocol^20^ after only one week of training, while all mice reached at least stage 6. In comparison, mice manually trained on the same protocol reached stage 2-3. When we investigated the learning curves as a function of trials completed, manually and automatically trained mice did not show a large difference. Therefore, the main reason for the difference in training efficiency was due to the higher number of trials mice completed during the 12 possible 15-minutes-long training sessions than during the single daily 30-minutes manual training, despite higher omission rate in the ATS, which could be a consequence of frequent access to water. Therefore, the automated training protocol may save significant amount of time otherwise spent by manually training the animals and, at the same time, results in better trained mice in substantially shorter time. Additionally, automated training does not require handling of mice, which is important in every manual training protocol to reduce animal stress caused by interaction with humans, thus saving the time otherwise spent on animal handling as well. Manual protocols often train animals with pellet rewards, daily sessions, and might involve a handling period before training so the animals get used to lab personnel. While scaling up manual training involves more human resources, increasing the number of mice trained in ATS (two mice per systems in parallel in the present implementation) only requires increasing the number of ATS setups. Since these systems are affordable and open source^32^, ATS provides a modular, readily scalable solution for mouse training. Future iterations may make use of RFID chips and decoders, thus allowing training multiple mice within the same ATS^12,33^.

Significant attempts have been made recently towards automated behavioral training^9,10,12,15,16,33–35^. One of the first automated training systems has been introduced in the Brody lab for training rats on a flexible and expandable set of decision making tasks^9,36,37^. It solved training with no human interaction and served as a prototype of later systems. Nevertheless, it did not provide a comparison with traditional training methods and thus it left the question open whether the hard-to-formalize experimental decisions during training such as when to advance between training stages, when to terminate a session, whether and when to introduce training breaks, etc. can be automated without a compromise in training efficiency. Another milestone was marked by the Olvecky lab that successfully combined automated training with automated recording in rats^8,10^, while the system was rather specific for that purpose. We have chosen the 5-choice serial reaction type task, a popular rodent paradigm^20,21,38–40^, that has also been the subject of previous automation studies^16,17^. We have built on these earlier works by both providing an affordable, flexible, modular system as well as a systematic comparison with manual training in terms of training efficiency.

It was shown that training animals on the same operant task using either food or water reward had similar mild effects on animal wellbeing, while animals receiving water reward acquired the task faster, and were more motivated to work for reward^41^. In addition, fluid reward avoids chewing artifacts, making it easier to combine with neuronal recordings; therefore, we modified the 5CSRTT protocol to provide water reward instead of food pellets, and demonstrated fast training with water rewards. Finally, we scaled up training speed by attaching two home cages to one training chamber and demonstrated that it is possible to train two mice simultaneously in an alternating fashion.

In experiments where uniform behavioral performance is important, it is beneficial that the animals receive ‘pre-training’ before they undergo virus injection or implantation surgeries^42,43^. The surgery often affects the performance of the animals, likely due to a combination of factors such as lack of training during the recovery period, changes in head dimensions altering the access to important spaces of the training setup due to the implants, the need of retraining muscles due to muscle trauma and altered balance, and increased stress^44–46^. Therefore, we separately tested the effect of surgeries and training breaks on 5CSRTT performance in the ATS. When we introduced a 17-days break after one week of training, we observed a transient decline in accuracy on the first day in the ATS-trained mice. This may be due to the more difficult task regime these animals experienced, as they resumed training at higher stages, according to their pre-pause levels, compared to manually trained mice. However, ATS-trained mice quickly regained performance, thus all training benefits of the ATS were maintained after the break. Similarly, transferring mice to the manual training setup after a second training break did not cancel the positive consequences of the ATS-training.

To further establish the assay’s practical use, we combined the automated training setup with wireless optogenetics^47^, broadening the range of possible experiments. Our design of two separate home cages connected with a single training chamber allows automatic training of mice that express the optogenetic actuator^48^ in parallel with control mice in the same training box, minimalizing the potential differences and uncontrolled factors between the two groups. It is important to remove potential subconscious biases in animal handling when performing optogenetic studies^2,4,5^, also achieved in this arrangement. Many freely behaving, trial-based, temporally controlled rodent task designs can be implemented in the behavior control system based on Bpod featuring five independent ports that can deliver either water reward or air-puff punishment with high temporal precision, by re-programming the open source finite state machine that controls transitions in the behavior protocol^1,33^. Since wireless optogenetics is controlled by TTL pulses synchronized with the behavior control, it is possible to precisely deliver photostimulation in any given task phase and part of the trial, allowing temporally specific manipulations^49–53^.

Animal stress may impede learning, increase behavioral and neuronal variability and therefore limit the interpretation of behavior neuroscience studies^54,55^. The increased variability may necessitate higher sample sizes, which, together with animal welfare concerns due to elevated stress, requires ethical considerations. We have partially eliminated important stressors during mouse training. Specifically, no human interaction was needed to carry out behavioral training in the ATS; additionally, mice were free to choose whether to engage in a given training session. Indeed, by measuring blood corticosterone levels, the main glucocorticoid stress hormone in rodents^25,26,28,29^, we found that training in the ATS caused significantly less stress to mice, which showed corticosterone levels similar to that of controls.

The Automated Training System provides a fully automated, experimenter-free training environment. The animals have the opportunity to train 12 times a day, which significantly speeds up learning. Participating in training sessions is not mandatory and the amount of water consumed depends on the individual animals’ thirst and willingness to perform, which lead to reduced stress in the training environment. Mice trained in the ATS system had no difficulty switching to manual training while retaining their performance levels. In the current implementation, two mice can be trained simultaneously on the 5CSRTT in one week, without any human interference. The system can readily be modified to train animals on a range of tasks, and we equipped the setup with wireless optogenetic stimulation to create an efficient, multi-purpose experimental tool.

## Methods

### Animals

Wild type male mice (N = 39, C57Bl/6J, over 6-weeks old) were used for the behavioral experiments and stress measurements; male homozygous ChAT-Cre mice (N = 7, over 2 months old) were used for surgical implantations and optogenetic experiments. All experiments were approved by the Committee for Scientific Ethics of Animal Research of the National Food Chain Safety Office (PE/EA/675-4/2016, PE/EA/864-7/2019) and were performed according to the guidelines of the institutional ethical code and the Hungarian Act of Animal Care and Experimentation (1998; XXVIII, section 243/1998, renewed in 40/2013) in accordance with the European Directive 86/609/CEE and modified according to the Directives 2010/63/EU. Food was provided ad libitum (Special Diets Services VRF1), while water access was scheduled as described in details below. A small, 15×5×2 cm 3D-printed box filled with nesting material served as nest in the ATS. All animals were kept on a 12-hour light-dark cycle. Light phase started at 7 am.

### Behavior setup

The ATS consisted of a central training chamber (16×16×10 cm) and two separate home cages, with controlled access to the training box. All chambers had grated floor with bedding underneath and were covered with a transparent plastic roof. Manual training was performed in an identical training chamber, but without the attached home cages. Manually trained animals were kept in standard mouse cages. The training chamber housed five adjacent water ports (Fig.1, 2A; Sanworks, US). Each port was equipped with an infrared photogate to measure port entry, a white LED to display visual cues, and tubing for water delivery connected to separate water containers for each port via fast, high precision, low noise solenoid valves (Lee Company, US). LED onsets, offsets and valve openings were controlled by printed circuit boards, connected to a Bpod open source behavior control system (Sanworks, US). The chambers were covered with soundproofing material^1^. A ‘house light’ LED was placed above the apparatus.

In the ATS, two 20×20×10 cm home cages were connected to the training chamber on each side through 10×5×4 cm tunnels. On both sides, the entrance to the training chamber was blocked by a motorized gate. The gates were equipped with infrared motion sensors (Panasonic EKMC series) attached to the roof of the home cage, directly above the tunnel entrance. Opening and closing of the gates was controlled by an Arduino Leonardo (Fig.1B-C). We set up a 24-hour surveillance system with web cameras and red lighting for the night period (Fig.1A-B). The cameras were accessed remotely to periodically check the operation of the ATS. Behavior control code was developed in Matlab and Arduino languages.

### Wireless optogenetic stimulation

The ATS was combined with a commercial wireless optogenetic stimulation system (NeuroLux, Fig. 1C). We wrapped the coil of the wireless system around the training chamber, which then created an electromagnetic field that powered an implanted micro-LED. The LED was emitting blue light (470 nm) upon induction through the coil. The optogenetic stimulation system allowed for precise, automated control of LED onsets an offsets by TTL signals^47^. Implanted mice were photostimulated during 50% of the inter-trial intervals in pseudorandomized order. Stimulation occurred at 20 Hz frequency and with 8 W.

### Training protocol

Mice were randomly assigned to two experimental groups. Water reward was used for motivation: animals undergoing manual training (N = 14) were subjected to a standard water restriction schedule, where they received water according to task performance during a 30 minute training session daily and additional free water for 2 hours/day, at least 2 hours after their last training session (from 2 to 4 pm). Animals trained in the ATS (N = 19) received their entire water intake from the task in the training chamber, accessed regularly every two hours for 15 minutes self-training sessions (Fig.2A-B). All ports of the training chamber delivered distilled water to avoid clogging of the tubing and valves; therefore, we placed a piece of mineral stone (Panzi, Hungary) as ion supplement in the home cages of the ATS. Weight of the animals was regularly monitored.

During 5-CSRTT, animals had to repeatedly detect flashes of light above one of the five ports presented in a pseudorandom order and report the detection by performing a nose poke in the respective water port. Upon correct reporting, 4-6 µl of water was delivered from the port as reward. Every session started with free access to 10-20 µl water from each port (in the manual training group, only in stage 1; Fig.2C). Each trial started with an inter trial interval (ITI), in which poking in the ports was prohibited. After the ITI, one of the ports was illuminated (Light On). The animal had to poke its snout into the illuminated port during ‘Light On’ or a short time period after that (limited hold, LH), in order to get the water reward. The length of the ITI, Light On and LH varied across training states as described by Bari et al.^20^. A poke during the ITI (premature response), in the incorrect port during Light On or LH (incorrect answer), or missing the periods allotted for nose poke (omission) resulted in a 5-second timeout, during which the house light was turned off. Each trial ended with either reward or a time-out punishment (Fig.2C).

We implemented a standard training strategy described by Bari et al.^20^. As detailed therein, the duration of the stimulus, ITI and LH was different from stage 1 to 12 to enable a progressive increase in difficulty. Mice were allowed to switch stages during a session in case they passed pre-defined criteria. Reward amount was set to 6 µl in stage 1, 5 µl in stage 2 and 4 µl in all subsequent stages. From stage 3, we randomized the duration of the ITI between 3, 4 or 5 seconds to increase attentional demand of the task.

### Surgery

Mice were anesthetized by an intraperitoneal injection of ketamine-xylazine mix (25mg/kg xylazine and 125mg/kg ketamine dissolved in 0.9% saline) and placed in a stereotaxic frame (Kopf Instruments, US). Local anesthetic (Lidocaine, Egis, Hungary) was applied subcutaneously and the eyes were protected by ophthalmic lubricant (Corneregel, Benu, Hungary). The skull was cleared and an opening was drilled above the horizontal diagonal band of Broca (HDB), a major hub of the central cholinergic system implicated in learning and attention^56,57^. A pipette pulled from borosilicate glass capillary was lowered into the target area and an adeno-associated viral vector (AAV 2/5. EF1a.Dio.eYFP.WPRE.hGH) was injected (300 nl to AP, + 0.75; MD, +/- 0.6; DV, - 5.0 and -4.7 mm). The wireless implant for optogenetics was lowered into the HDB (AP, + 0.75; MD, +/- 1; DV, - 5.5 mm). We secured the ring-shaped optogenetic sensing module to the surface of the skull with tissue glue (Vetbond, 3M, US). The needle that held the LED was cemented to the skull with dental cement (Paladur, Dentaltix, Italy). The skin above the implant was sutured and antibiotic cream (Baneocin, Medigen, Hungary) was applied on to the surgical wound. The animal was placed on a heating pad for recovery. A 2-weeks rest period was allowed for full recovery, after which the experimental protocols were initiated.

### Measuring the stress level of the animals

To measure acute stress of the animals caused by training (and handling in the case of manually trained animals), blood samples were collected after their last training session. On the 7^th^ day (9am to 12pm), manually trained animals were placed in their home cages after training for 10 minutes. After training at matching time of the day, the animals in the ATS were allowed to return to their home cages within the system for 10 minutes. Water consumption was similar in the two groups during the last training sessions. After the 10 minutes rest, mice were transferred to a separate room. For corticosterone level measurements, blood samples were collected during decapitation in ice-cold plastic tubes, centrifuged and the serum was separated and stored at -20 °C until analysis. Corticosterone was measured in 10 μl unextracted serum or undiluted medium by a radioimmunoassay (RIA) using a specific antibody developed in our institute as described earlier^58,59^. Samples from each experiment were measured in a single RIA (intra-assay coefficient of variation, 7.5%). We compared data after one week of training in three groups (control, N = 6; manually trained, N = 6; ATS-trained, N = 6). In the control group (N = 6), mice had food and water available ad libitum and were not handled. The behavioral data of these animals were included in Figures 3-5.

### Statistics

Behavioral performance was analyzed by custom-written open source code in Matlab 2016b (MathWorks, US) available at https://github.com/hangyabalazs/ATS. Statistical analysis was carried out using the STATISTICA 13.4 software (TIBCO, US). Group differences were assessed by one-way, repeated measures ANOVA. Newman-Keuls post hoc tests were performed after ANOVA if the main effects were significant. Wilcoxon singed rank test and Mann-Whitney U-test were used for non-parametric comparison of central tendencies between two paired or unpaired distributions, respectively. Data are presented in the figures as mean ± standard error. Differences were considered significant at p < 0.05.

## Data availability statement

The datasets generated and/or analysed during the current study are available from the corresponding author on reasonable request. Code for hardware control and behavioural data analysis can be downloaded from https://github.com/sanworks/Pipeline_Gate and https://github.com/hangyabalazs/ATS).

## Acknowledgements

We thank Katalin Lengyel and Éva Dobozi (Semmelweis University, Budapest, Hungary) for expert technical assistance in anatomical methods and hormone measurements; Drs. Éva Mikics and László Acsády for helpful comments on the manuscript. This work was supported by the ‘Lendület’ Program of the Hungarian Academy of Sciences (LP2015-2/2015), NKFIH KH125294 and the European Research Council Starting Grant no. 715043 to BH. BH is a member of the FENS-Kavli Network of Excellence.

## Author contributions

EB, DB and BH conceived the project, EB and DB performed the experiments with continuous engineering support from JIS, AB performed stress hormone measurements, DB and BH supervised the project, EB, DB and BH wrote the manuscript with input from AB and JIS.

